# Differential kinematic control and co-ordination among redundant joints during whole arm reaching movements

**DOI:** 10.1101/2025.10.08.681178

**Authors:** Niranjan Chakrabhavi, Ashitava Ghosal, Aditya Murthy

**Author notes:** **Corresponding Authors** – Niranjan Chakrabhavi and Aditya Murthy. **Author Contribution:** NC conceived the study; NC and AM designed the experiment; NC collected data. NC performed the data analysis; AM and AG supervised the analysis; NC and AM wrote the manuscript; NC, AM, and AG, reviewed the manuscript. **First Author:** Niranjan Chakrabhavi.

## Abstract

Normative upper limb movements are produced by multiple redundant joints. While the reaching task is specified at the endpoint, such task objectives become implicit at the level of joints. A fundamental question is whether planning and control of joints is solely in the service of the endpoint or whether they also include joint trajectories. Using Spearman’s correlation and zero crossings, we found differential kinematic signatures of control between shoulder and elbow joints in contrast to the wrist joint. However, the extent of control among joints was substantially diminished compared to the endpoint. Further, when such control measures were compared to the subspaces of inter-trial joint exploration, we found that online control at proximal joints, such as the shoulder and elbow, were significantly associated in regulating the task space, while control at the wrist (distal) joint was associated in regulating joint redundancy in null space. These results suggest that null space is not entirely uncontrolled as per the uncontrolled manifold hypothesis but selectively controlled by some distal joints. Additionally, across different directions, either the shoulder or the elbow contributed dominantly towards the movement of the endpoint while the other joint was lagging and that this strategy reflected in our kinematic measures of online and trajectory control. Taken together, this study shows how the selective implementation of a leading joint in task space and a lagging joint in null space can enable the control of multi-jointed movements and attenuate the problem of joint redundancy.

**Significance Statement:** In this study, we addressed a fundamental question of how our central nervous system resolves the problem of joint redundancy, as to whether planning and control of joints are solely in the service of endpoint or whether they also include joint trajectories. We found significant kinematic control signatures among joints towards their respective average joint trajectories, especially during the early and middle phases of movement. Furthermore, each joint had a distinct task objective, the proximal shoulder or elbow joints controlled the task space and were responsible for driving the whole-arm movements, while the distal wrist joint regulated joint redundancy.

## Introduction

In human beings, behavior largely manifests with the production of multi-joint movements. This involves controlling individual joints as well as spatiotemporal coordination between them (1, 2). Common daily activities such as reaching for an object, walking, typing on a keyboard, and so on, utilize multi-joint movements. These seemingly easy tasks are computationally challenging for our central nervous system (CNS). For example specifically, our upper arm is bestowed with seven degrees of freedom (DoFs), while reaching for targets in cartesian space poses only three constraints i.e., the location of the target (3). The challenge for the CNS is to solve this under-constrained problem to arrive at a single solution for the trajectories of joints and endpoint (hand) among infinitely many possible theoretical solutions, also known as the problem of motor redundancy (4).

One way to address this problem of redundancy or abundancy as it is sometimes referred to (5), is to look at signatures of planning and control among joints relative to the endpoint. Numerous studies on reaching have shown relatively stereotypic straight-line trajectories with bell-shaped velocity profiles at the endpoint. In contrast, the pattern of joint movements have been shown to be less stereotypical and are accompanied by greater inter-trial and inter-task variability (4, 6). Such observations have inspired the formulation of the uncontrolled manifold hypothesis (UCM), which was developed (5, 7, 8) to mitigate the problem of redundancy by investigating the degree of coordination among the joints (4, 9). In the UCM framework, the inter-trial joint variability is projected onto two subspaces, the task space and the null space. According to the hypothesis, the joint projection onto the task space affects the endpoint and is hence required to be tightly controlled, while the projection onto the null space does not change the endpoint, also named as the uncontrolled manifold, where the CNS does not enforce control over the variability along this manifold (10). The UCM hypothesis is also closely related to the minimal intervention principle (11) where controlling solely the task space is considered an optimal strategy and together provide a coherent explanation of normative behaviour in a wide range of tasks involving redundant motor systems (1).Thus, the production of movements through control of individual joints, has been largely ignored in the study of human reach movements.

In contrast to the UCM hypothesis, the leading joint hypothesis (LJH) proposes a mechanism of resolving redundancy by explicit control over fewer joints (2, 12–17). According to LJH, the brain chooses one or more leading joints and produces active muscle torque to move the serially linked arm. Whereas, the other subordinate/lagging joints utilize interaction torques for their movements and are monitored constantly to stabilize themselves and to achieve endpoint accuracy during the movement (2, 13). In the context of upper limb movements, it has been observed that the leading joint is generally the proximal shoulder joint, but it switches over to the elbow joint in a few cases (16, 17). According to LJH, the approach of joint control is in the framework of limb kinetics, as a mutual competition between the active muscle torque and the interaction torque at any joint. Further, the contribution of individual joint rotations in producing the endpoint velocity is utilized as a kinematic proxy to identify the leading and the lagging joints (12, 13). Despite that, it is unclear whether there exists a similar representation of kinematic control among joint trajectories, i.e., whether some joint trajectories, such as the trajectories of the leading joint, undergo more kinematic control/corrections compared to the other joints and whether the kinetic and the kinematic approaches of joint control correspond with each other.

Multiple attempts have been made to quantify the extent of online control during various motor tasks, often relying on linear regression, as the lack of predictability of end-of-movement characteristics like movement amplitude and movement time by early kinematic markers such as peak velocity and peak acceleration (18–21). To find earlier signatures of trajectory correction, the postures during different phases of movement have been correlated to the postures at the end-of-movement (22–24), largely focused on the analysis of endpoint trajectories. These techniques, however, are limited to linear associations, susceptible to changes in intra-trial variability and are sensitive to outliers (25). In this work, we resolved the above limitations by attributing crossovers among trajectories as a robust signature of online control and detected them using Spearman’s rank correlation. Furthermore, instead of correlating postures referenced to the end-of-movement, we probed for rapid signatures of online corrections by correlating the rank order of trajectories at movement onset with the respective rank order at subsequent phases of movement (26).

However, it was still unclear whether such corrections were directed towards a desired trajectory (27, 28). So, we developed zero crossings rate (*ZCR*) to detect crossovers of trajectories across a trial-averaged trajectory, to estimate the extent of real-time corrections implemented by the CNS to follow an average trajectory. We also probed the extent of kinematic trajectory control during early, middle and late phases of movement.

Further, we compared the extent of kinematic control, using the above two measures, among the movement trajectories of the three joints and the endpoint. In the context of LJH, we used the velocity projection approach to identify the leading and the lagging joints (12, 13), and hypothesized that the trajectories of the leading joint undergo greater kinematic control than the lagging joint. Furthermore, we tested whether the notions of joint control (LJH) and joint coordination (UCM) can be reconciled to explain how multi-jointed reaching movements are planned and controlled. In the context of UCM hypothesis, any online correction at joints is posited to be associated with regulating the task space (or the endpoint) and not the null space. We tested this by systematically comparing our measure of kinematic online control at individual joints with the overall joint projections onto the task space and the null space, respectively.

## Materials and Methods

This study involved upper limb horizontal reaching movements performed by 15 right-handed participants (Age: 21-32) and were selected using Edinburgh Handedness Inventory (29). The experiment was approved by the institute ethics committee at IISc Bengaluru, India, and participants provided their informed consent before they started the experiment.

### Experimental Setup

3D electromagnetic motion trackers (Teardrop Mini sensors, 240Hz, Liberty tracking system, Polhemus Inc., Colchester, USA) were secured over the skin at the right sternoclavicular joint (neck), right acromion process (right shoulder), lateral epicondyle (elbow), dorsal tubercle of radius (wrist) and the dorsal aspect of the hand (endpoint) with 3M micropore surgical tape. The subjects held on to an endpoint robotic manipulandum (KINARM, 240 Hz, BKIN Technologies Ltd., Kingston, Canada) which was mapped to a pink cursor on monitor screen in front of them (Fig 1A). The experiments were programmed on TEMPO/VideoSYNC Software (Reflective Computing, St. Louis, MO, USA). The data was processed and visualized on MATLAB interface (MathWorks, Massachusetts, USA).

**Figure 1:**
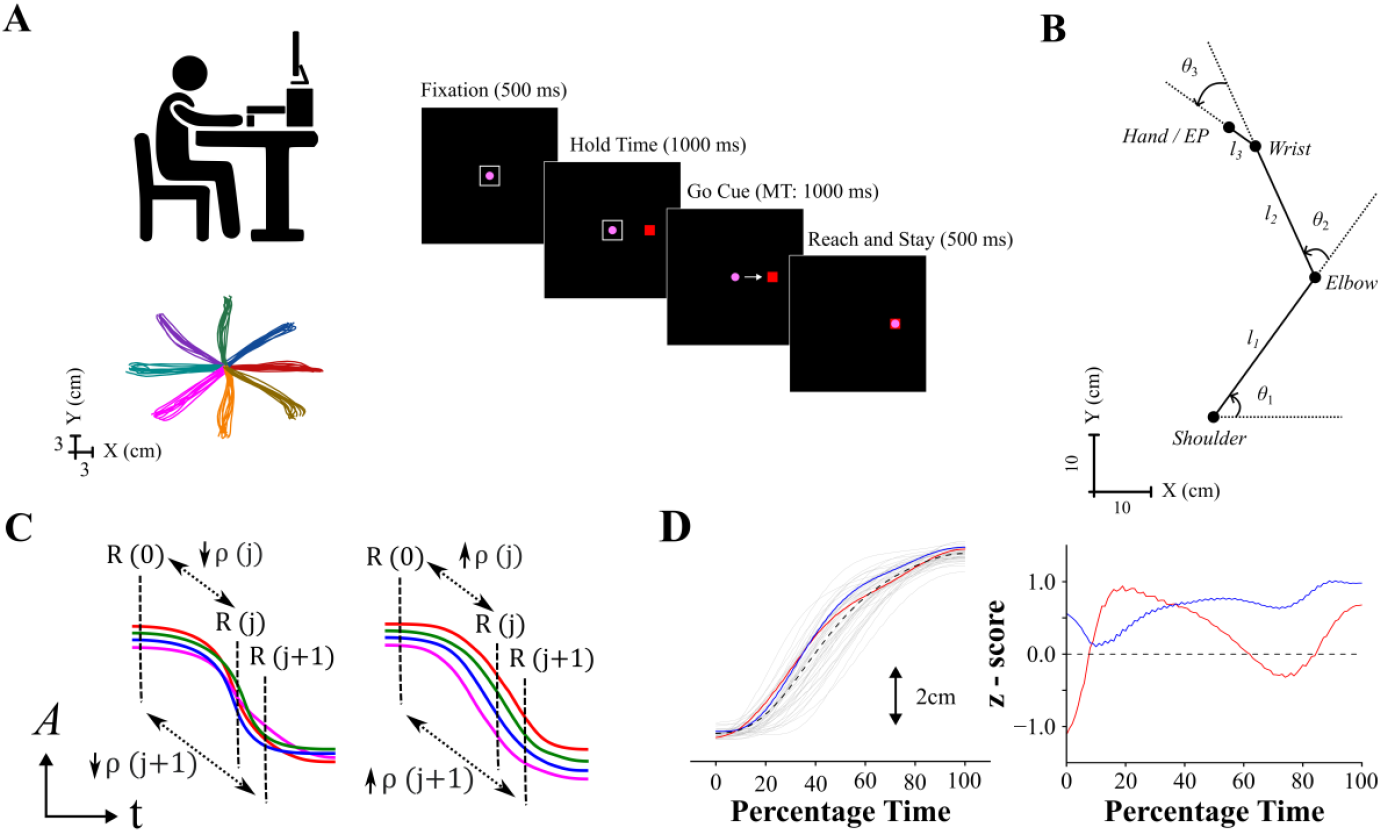
Experimental design, the kinematic model of the upper limb and the idea of online and trajectory control. A) The experimental apparatus, the experimental protocol and sample trajectories (10 trials in each direction) traced by the hand/endpoint (EP) of a single participant. B) The upper limb is modelled as a 3-link serial chain consisting of shoulder, elbow, and wrist joints, with one rotational degree of freedom at each joint, and subtending angles θ_1_, θ_2_, and θ_3_ respectively. The upper arm, the forearm and the palm had link lengths of l_1_, l_2_, and l_3_ respectively, as shown in the figure. MT – movement time. C) Conceptual representation of online control (left): Consider initial rank order R(0), with four representative trials marked by red, green, blue, and pink lines, such that any online correction thereafter of trajectories shall lead to crossovers among them (rank order R(j+1), at time t=j+1, becomes pink, green, red, and blue) and can be captured by performing the Spearman’s rank correlation between the rank order of initial posture and subsequent postures, from time t = 0,…,j,j+1,…, during the movement to obtain a time varying ρ(t). The extent of decorrelation among trajectories is an indicator of online control or corrections. However, if there are no crossovers observed among trajectories (right), it would be attributed to weak/lack of control. A-amplitude, t-time (percentage time between movement onset and movement end or on actual time scale with first 500 ms post movement onset). D) Sample trajectories of X coordinate of endpoint plotted against percentage time during movements along 0^0^ direction (left). The dashed black line is the average trajectory, and the blue and red lines are two demonstrative trajectories. The trajectory locations are projected onto their respective z – scores (right, see methods). It can notice that the red line crosses the average trajectory three times during the movement and is considered to be in the influence of greater trajectory control while the blue line never crosses over and hence is considered to be under weaker trajectory control.

### Experimental Protocol

The participants fixated at the center for 500 ms, upon which a red colored target appeared in the periphery at 12 cm eccentricity. The location of the target was block randomized at any of the 8 directions separated by 45º (Target locations - 0º, 45º, 90º, 135º, 180º, 225º, 270º, and 315º). After a delay (hold time) of 1000 ms, the go cue was presented by the disappearance of the fixation box. Subjects had to initiate a simple center out reaching movement towards the target location within a movement time (MT) of 1000 ms and stay there for 500 ms to complete one trial. The inter-trial interval was 1500 ms and subjects performed ∼40 trials (5 out of 15 subjects performed ∼60 trials towards 0^0^ and 45^0^, and ∼20 trials towards other directions) along each of the eight directions (sample trajectories are shown in Fig. 1A). The participants subsequently performed the same task with the contralateral upper limb, and the sequence of dominant and non-dominant arms were randomized. The data from the dominant arm has been chosen for further analysis.

### Data pre-processing

The motion data from the sensors were filtered using a fourth-order, zero-lag Butterworth filter at a cutoff frequency of 20 Hz (30). Movement onset and end were identified by detecting when the hand crossed 5% of maximum speed (31–33) and then descended till the speed reached zero for the first time. The motion data were normalized into 100 equally spaced bins for the analysis involving percentage time scales and the first 500 ms was used to detect the timing of control. The analysis of fixation condition was performed on the data 1000 ms prior to movement onset.

#### Model of the upper limb and calculation of joint angles and link lengths

The upper arm was approximated as a 3-link model involving rotations at the shoulder, elbow and wrist joints (Fig. 1B). The participants were required to make simple horizontal reaching movements between the fixation and the target location, while the orientation of the hand holding the robotic manipulandum was not constrained. Hence the task posed two constraints, i.e., the location of the target in the XY plane, whereas these movements were produced by rotations at three joints – constituting to the three degrees of freedom. The system could choose among infinitely many redundant trajectories between the fixation and the target; and having chosen a particular trajectory, the arm would still have one redundant degree of freedom at the joints.

The angular rotations at the shoulder, elbow, and wrist joints (Fig. 1B) were calculated by the angle subtended between two vectors around each of the joints using the tan2d function (MATLAB, MathWorks, Massachusetts, USA). The angle between a vector connecting the shoulder to elbow, with a unit vector along the X axis, was used to track horizontal adduction– abduction movement at the shoulder joint (*θ*_1_). The angle subtended between a vector connecting shoulder to elbow, with a vector connecting elbow to wrist, was used to measure the flexion - extension movements at elbow joint (*θ*_2_) and similarly for the flexion – extension movements at the wrist joint (*θ*_3_), as depicted in Fig. 1B.

The link lengths of the upper arm (*l*_1_), forearm (*l*_2_), and palm (*l*_3_), were calculated using the Euclidean distance between the locations of sensors (in the XY horizontal plane) around each of the joints and were averaged across the duration of the experimental session.

Having computed the three joint angles *θ*_1_, *θ*_2_, and *θ*_3_, and the three link lengths *l*_1_, *l*_2_, and *l*_3_, constituting the upper arm, a forward kinematic model was constructed to predict the endpoint (hand) movements in the XY plane using Eq. 1. The model predictions were then compared with the actual measured movements of the endpoint using Pearson’s correlation coefficient.

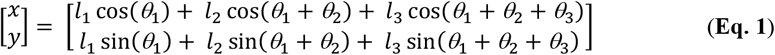

where *x* and *y* correspond to the endpoint locations in the X and Y directions, respectively.

#### Control of joints through decorrelation and zero crossings

The time series data of the three joint angles and the endpoint (in the direction of X and Y) were first subjected to Canonical Correlation Analysis (CCA) as described in Krüger et al., 2017 (24), but in this case, between the starting postures and their respective postures at subsequent time instances, to compare online control at the level of joints relative to control of hand/endpoint (Fig. 2A).

**Figure 2:**
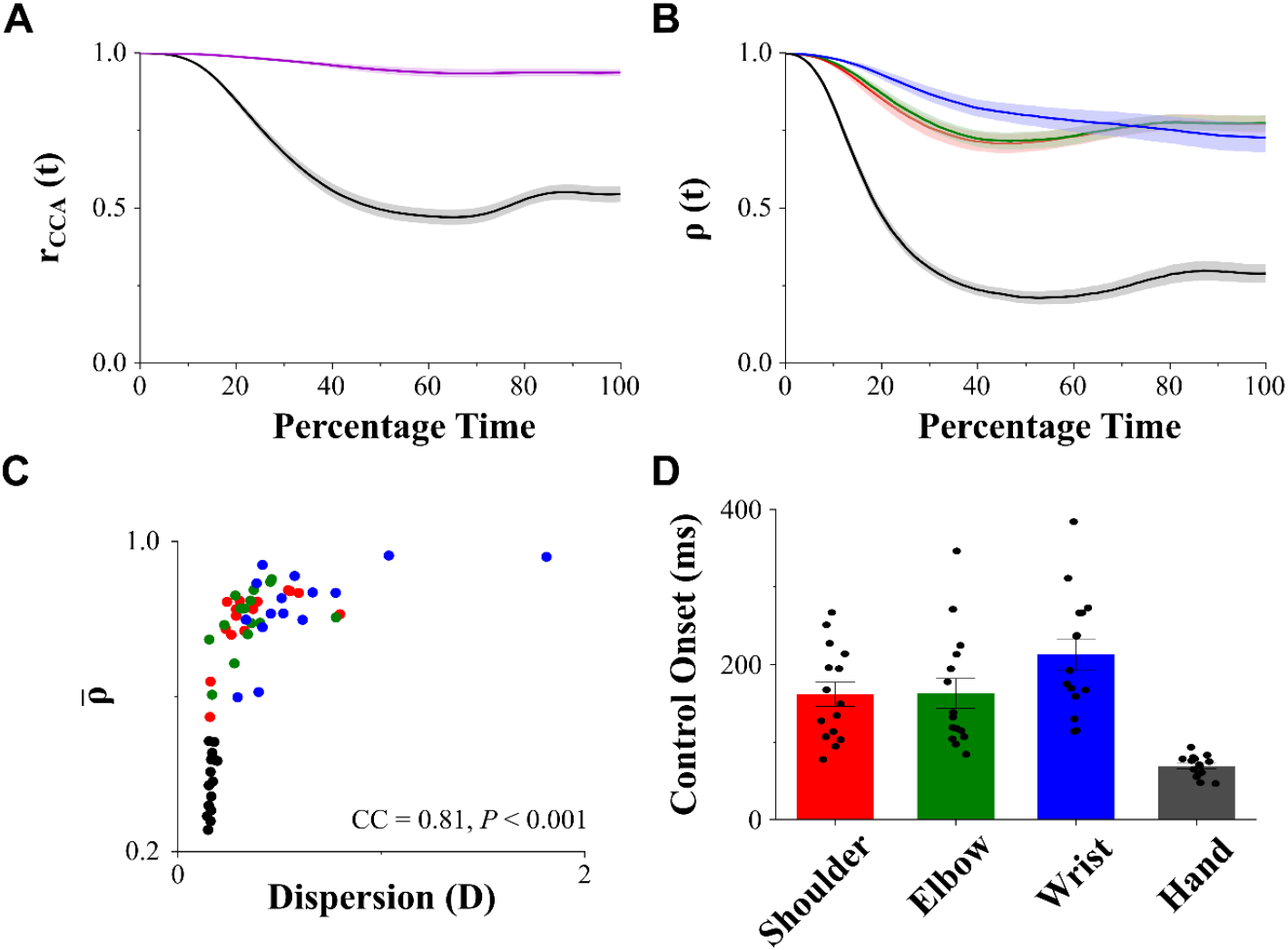
Signatures of online control among joints and endpoint (hand). A) Canonical correlation analysis (CCA, mean ± SEM) performed between the starting and subsequent postures on trajectories of joints and endpoint. Decorrelation was greater in the trajectories of endpoint when compared to that among joints. B) Spearman’s correlation analysis (mean ± SEM) performed on the shoulder, elbow, wrist, and endpoint. Results were qualitatively similar to that in A. C) Association between average decorrelation measure with the average trajectory dispersion among the joints and endpoint. We found significant association between them (Spearman’ correlation co-efficient, CC = 0.81, P < 0.001). D) The onset of control (using the tangent method, mean ± SEM) of joints and endpoint. We found significantly delayed initiation of online control among joint trajectories when compared to the endpoint (Adj. P < 0.001). Among joints online control of wrist was delayed with respect to shoulder (Adj. P = 0.043) and elbow (Adj. P = 0.05) joints. The colors red, green, blue, and black represent shoulder, elbow, wrist and hand (endpoint) respectively. The purple colour in A represents joints.

Even though we could compare the extent of control between joints and endpoint using the above analysis, distinguishing control among the various participating joints was not possible. Also, we know that the above analysis and those described previously in literature are sensitive to changes in inter-trial variability and are sensitive to outliers (25). We also know from the literature (22, 24) that the inter-trial variability varies during the time course of movement. Therefore, we used Spearman’s rank correlation to capture crossovers among trajectories by comparing the rank order at the start of movement, *R*(0), with the rank order at every subsequent time-instance, *R*(*j*), where *j* ranged between movement onset to end of movement (0 to 100%) for analysis on percentage time scale, and between 0 to 500 ms for analysis on actual timescale. In Fig. 1C, the left panel depicts greater online control with significant crossovers among the four representative trajectories leading to a decrease in Spearman’s rank correlation (*ρ*(*t*)) over the time course of movement, while the right panel shows weaker/lack of online control with the absence of such crossovers (*ρ*(*t*) equal to or closer to 1). We associated the lack of correlation (decorrelation), i.e., the correlation co-efficient averaged across the percentage time stamps 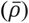, to the extent of online control (Figs. 2, 4, 5, and S2). We went ahead and fit a logistic function, *ρ*_*f*_(*t*), to the time series of correlation co-efficient, *ρ*(*t*), for each of the subjects and used the value, *β*, to quantify the rate of decorrelation or crossovers as per Eq. 1, where, *β, a* and c, were free parameters.

**Figure 3:**
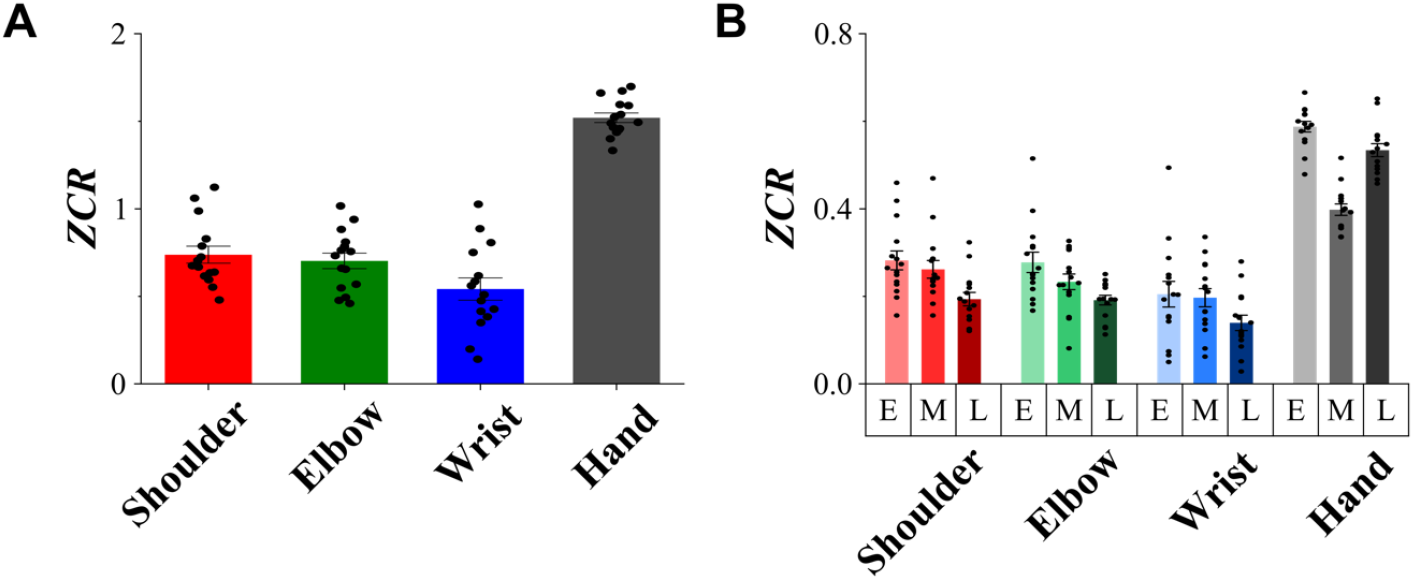
Zero Crossings Rate (ZCR) of joints and hand (endpoint). A) ZCR (mean ± SEM) of shoulder, elbow, wrist, and endpoint. We found significantly higher zero crossings at the endpoint with respect to the joints (P < 0.001). B) ZCR (mean ± SEM) of joints and endpoint computed during early (0 to 33 %, light), middle (34 to 66 %, medium), and late (67 to 100 %, dark). We found higher zero crossings during early and late phases at the endpoint, but it was higher only during the early phase at the joints. Colours: shoulder – red, elbow – green, wrist – blue, and hand (endpoint) – black.

**Figure 4:**
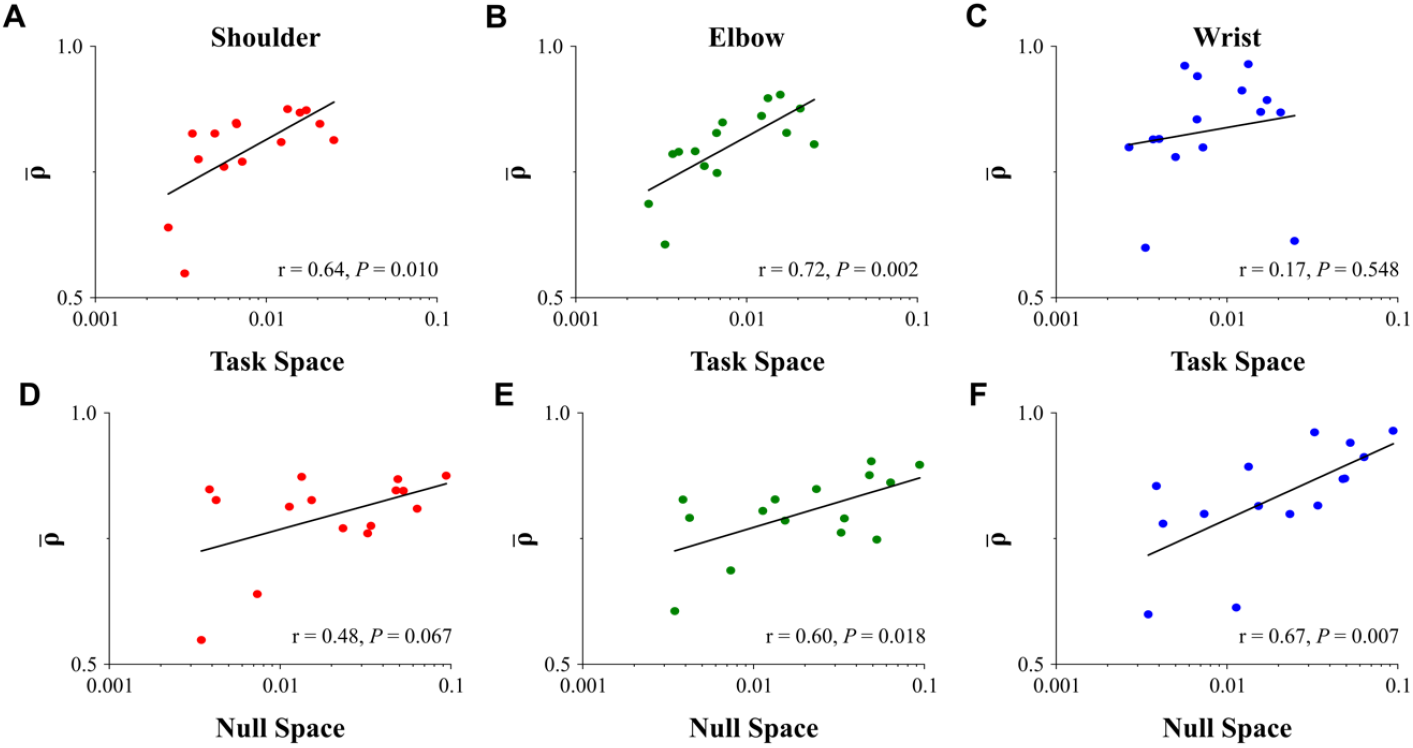
Association of the average decorrelation measure with task space (top) and null space (bottom) exploration of the shoulder (red), elbow (green), and wrist (blue) joints. We found that the average decorrelation of shoulder (A) and elbow (B) were significantly associated with the exploration in the task space, but it was not the case for the wrist joint (C). We also found that the average decorrelation of elbow (E) and wrist (F) were significantly associated with the exploration in the null space, but it was not significant for the shoulder joint (D). The horizontal axes were logarithmically scaled with base 10.

**Figure 5:**
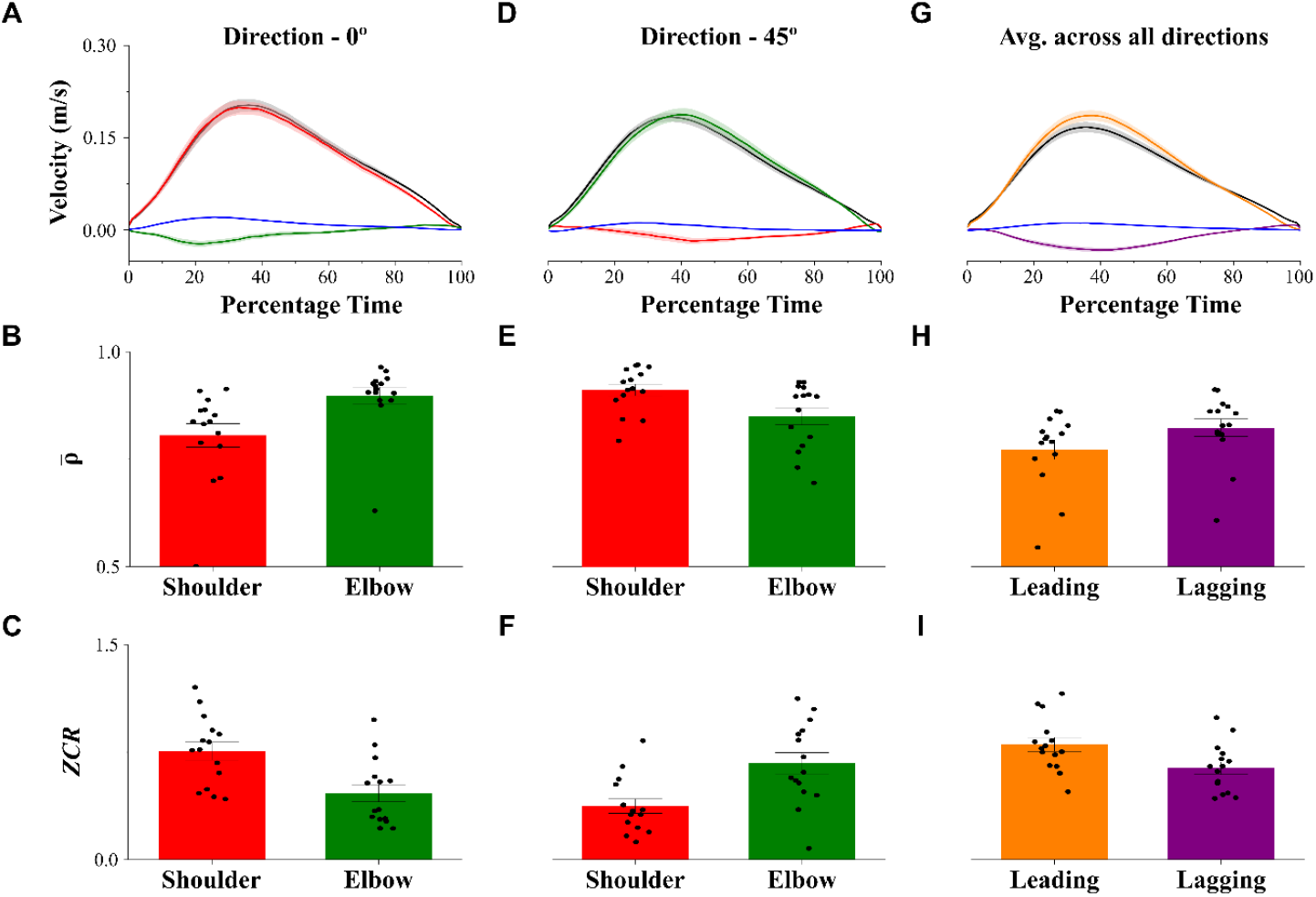
Leading joint hypothesis, average decorrelation and zero crossings rate. All quantities plotted here – mean ± SEM. Colors: red – shoulder, green – elbow, blue – wrist, black – hand (endpoint), orange – leading joint, purple – lagging joint. For movements towards 0º, A) shoulder contributed to most of endpoint velocity. B) The shoulder had significantly greater decorrelation than the elbow (P < 0.001) and C) ZCR at shoulder was significantly greater than the elbow (P < 0.001). For movements towards 45º, D) elbow joint contributed to most of endpoint velocity. E) The decorrelation at elbow was significantly greater than that at the shoulder (P < 0.001) and F) the ZCR at elbow was significantly greater than the shoulder (P < 0.001). Averaging the contribution of leading and lagging joints across all the 8 directions of movement. G) Leading joint contributed to most of the endpoint velocity. Wrist contributed to less than 6% of endpoint velocity overall (P = 0.02). H) The decorrelation at the leading joint was significantly greater than the lagging joint (P < 0.001) and I) the ZCR was significantly greater at the leading joint than the lagging joint (P < 0.001).

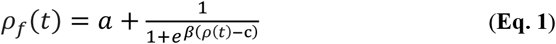

Further, to assess whether such crossovers are beneficial to the task, we associated the time-averaged decorrelation measure 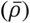, with the trajectory dispersion (*D*, Eq. 2), which was quantified as the ratio of average inter-trial standard deviation to the average range of movement during the *n* trials. The rationale was to evaluate whether a greater extent of trajectory crossovers would result in decreased trajectory dispersion. The measure was unitless and normalized for differences in movement amplitudes across joints and endpoint.

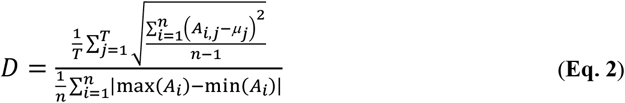

where, *A*_*n x T*_ was the amplitude matrix with *n* trials across *T* = 101 percentage time steps between movement onset and movement end, *μ*_*j*_ was the average inter-trial amplitude at time *j*, and *A*_*i*_ was a single trajectory on the *i*^*th*^ trial.

The onset of online control was detected using the “tangent method”, a technique earlier used to find the end of lag phase and initiation of growth phase in a growth curve (34, 35). At first, the Spearman correlation co-efficient (*ρ*(*t*)) was found for the first 500 ms into movement execution as described in Fig. 1.C. A tangent was constructed at the point of maximum rate of decorrelation, and the onset of online control was marked as the time at which the tangent intersected the *ρ* = 1 horizontal line.

We were further interested whether trajectory corrections during movement were towards an average planned trajectory. We did this by computing *z* – scores (*zi,j*) for every *i*^th^ trial and at every percentage time stamp *j*, using Eq. 3.

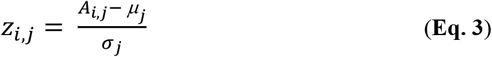

where, *σ*_*j*_ – inter-trial standard deviation at time stamp *j*.

Sample reach trajectories (grey), performed by a single participant towards 0º oriented target, are shown in Fig. 1D (left panel), where the dashed black line represents the inter-trial average trajectory which transforms to *z* = 0 line (Fig. 1D, right). The extent of corrections towards the average trajectory (Fig. 3A) was estimated by counting the number of times the trajectories crossed over the *z* = 0 line for a given number of trials, which we called the zero crossings rate (*ZCR*) as described in Eq. 4. The red and blue traces are two representative trajectories (Fig. 1D, right); while the red trace crosses over the *z* = 0 line thrice indicating a strong propensity towards the average trajectory, the blue trace does not show any crossovers and is said to be under weak/lack of trajectory control.

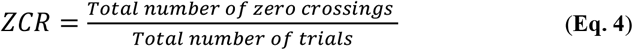

Subsequently, we also quantified the zero crossings rate during the early (0 – 33 %), middle (34 – 66 %) and late (67 – 100 %) phases of movement to investigate whether the pattern of control is constant or if it changes in unique ways for movements of joints and endpoint (Fig. 3B).

#### Joint coordination and quantification of redundancy

Joint coordination (Figs. 4 and S1) was analyzed by computing the null space and the task space projections of deviations in joint configuration (30, 36). From Eq. 1, changes in endpoint location were related to changes in joint configuration using a Jacobian matrix *J* with size 2 × 3, in Eq. 5.

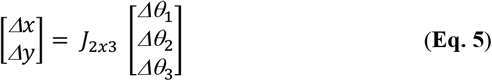

The Jacobian matrix (refer SI Methods for the expansion of *J*) was expressed as follows in Eq. 6 –

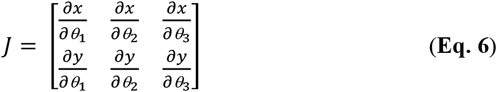

For a given trial *i*, and at every percentage time point *t*, the inter-trial deviation in joint configuration (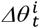, with size 3 × 1) was computed by subtracting the trial-averaged joint configuration 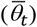 from the current joint configuration 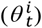 in Eq. 7.

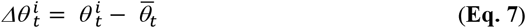

The null space vector (*ξ*) of the Jacobian matrix at the referent average joint configuration 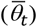 was found using Eq. 8.

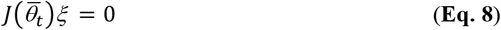

The changes in joint configuration, which did not affect the endpoint movements, the null space projection 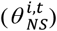, was found by projecting the joint deviation over the null space vector (*ξ*) of the Jacobian matrix in Eq. 9.

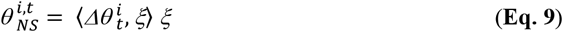

Further, the changes in joint configuration which resulted in movements of the endpoint, the task space projection 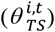 was computed using Eq. 10.

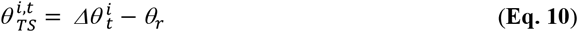

The extent of redundancy or the null space (*NS*) variability was defined as the averaged sum of squares of the null space projection across *n* trials and through the percentage time from 0 to 100 % in Eq. 11.

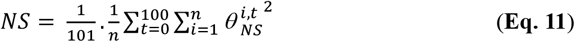

The extent of task space (*TS*) variability was defined as the averaged sum of squares of the task space projection across *n* trials and through the percentage time from 0 to 100 % using Eq. 12.

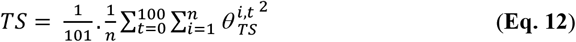

#### Velocity projection of joints on endpoint

We computed the angular velocities (***ω***_*k*_) of shoulder (*k* = 1), elbow (*k* = 2) and wrist (*k* = 3) joints and related them to the velocity of the hand/endpoint (***V***_*h*_) using Eq. 13 (12, 13).

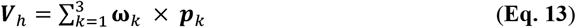

Where ***p***_*k*_ is the position vector between *k*^th^ joint and the endpoint.

The contribution of each joint (*V*_*k*_) towards the endpoint velocity was calculated by taking the dot product with the unit vector along the endpoint velocity 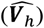 using Eq. 14. The ratio of the velocity contribution of individual joints (*V*_*k*_) with the magnitude of endpoint velocity (*V*_*h*_) was calculated at each percentage timestamp, was multiplied with 100, and the median of this quantity across timestamps, was used to evaluate the leading and the lagging joints (Figs. 5 and S2). We settled with the kinematic proxy of LJH and did not go ahead with the torque analysis as the double derivative of joint trajectories were noisy.

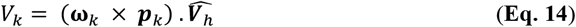

To test whether the joint leading the movement was also associated with an enhanced and a faster degree of online control, we compared the average decorrelation measure between the leading and lagging joints. We also evaluated whether the leading joint followed a planned trajectory compared to the lagging joint by comparing their *ZCR* measures.

The results through Figs. 2-4, 6 and S1 were averaged across all movement directions while individual directions were explored in Figs. 5 and S2. The decorrelation and dispersion measures were further averaged across the X and Y dimensions for the analysis of endpoint.

#### Statistics

We performed a pairwise t-test for paired comparisons and a two-sample *t*-test for independent comparisons. For more than two sample paired comparisons, we performed repeated measures ANOVA. Multiple pairwise comparisons were reported with post-hoc Tukey test to limit Type – 1 errors.

## Results

The link lengths of the upper arm, forearm, and wrist and the corresponding joint angles at the shoulder, elbow, and wrist were calculated from the motion data. They were used to predict the endpoint trajectories in the X and Y directions using the forward model. The actual endpoint trajectories closely matched with the estimated endpoint trajectories, with a median [Q1, Q3] explained variance of 88.31 [83.7, 90.9] % across subjects (Q1 – first quantile, Q3 – third quantile).

### Online control of joints and endpoint during reaching movements

We wanted to compare the extent of online control among the joints to that at the endpoint. At first, we subjected the trajectories of joint angles and the endpoint to canonical correlation analysis (Fig. 2A). We found that the average correlation co-efficient, the proxy for online control, was significantly less at the endpoint (mean ± SEM: 0.64 ± 0.02) as compared to that among the joints (mean ± SEM: 0.96 ± 0.01, *P* < 0.001), indicating that the online control at the endpoint was stronger than at the level of joints.

To distinguish control among the different joints and endpoint (Fig. 2B), we subjected the trajectories to Spearman’s correlation analysis. We performed a 1-way repeated measures ANOVA on the average decorrelation (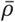, levels: 4 - shoulder, elbow, wrist, and endpoint) showed a significant difference between them (*P* < 0.001). Further, the pairwise Tukey test revealed that the average correlation coefficient of the shoulder (mean ± SEM: 0.80 ± 0.02), elbow (0.80 ± 0.02), and wrist (0.83 ± 0.03) joints were significantly greater than that of the endpoint (0.38 ± 0.02, Adj. *P* < 0.001 each) suggesting that the online control of endpoint was much higher than among the joints. We performed a similar 1-way repeated measures ANOVA on the rate of decorrelation (*β*) and found a significant difference among the four groups (*P* < 0.001). A pairwise Tukey test showed substantial differences between endpoint (mean ± SEM: 0.143 ± 0.005), and shoulder (0.078 ± 0.007, Adj. *P* < 0.001), elbow (0.074 ± 0.004, Adj. *P* < 0.001), and wrist (0.036 ± 0.005, Adj. *P* < 0.001) joints, respectively. We also found significant differences between shoulder and wrist (Adj. *P* < 0.001) and elbow and wrist (Adj. *P* < 0.001) joints, but not between shoulder and elbow joints (Adj. *P* = 0.92) despite a greater exertion of the elbow joint (range, mean ± SEM: 18.2º ± 0.5º) as compared to the shoulder joint (12.0º ± 0.3º, *P* < 0.001).

Further, we investigated whether the joint decorrelation was meaningful or merely due to noise by performing three analyses. Firstly, we found that the average decorrelation in joints was significantly associated with joint dispersion (Fig. 2C, Spearman’s *ρ* = 0.62, *P* < 0.001). When the endpoint was taken into consideration along with joints, the association became stronger (*ρ* = 0.81, *P* < 0.001). Next, we performed 1-way repeated measures ANOVA (levels – 8 directions of movement) on the average decorrelation among joints, and we found that it was sensitive to movement direction for shoulder (*P* < 0.001), elbow (*P* < 0.001) and wrist (*P* = 0.01) joints. Finally, we compared the average decorrelation during goal-directed movement of the joints to that during fixation at the center by performing a 2-way repeated measures ANOVA with two factors (goal and fixation) and three levels (shoulder, elbow, and wrist joints) and found a significant difference between goal and fixation conditions (*P* < 0.001). Pairwise comparisons showed significantly greater decorrelation during goal movements for shoulder (mean ± SEM, goal: 0.80 ± 0.02, fixation: 0.99 ± 0.004, Adj. *P* < 0.001), elbow (goal: 0.80 ± 0.02, fixation: 0.99 ± 0.002, Adj. *P* < 0.001), and wrist (goal: 0.83 ± 0.03, fixation: 0.99 ± 0.001, Adj. *P* < 0.001) joints.

To further establish the distinction between endpoint and joint level control, we quantified the differences in the onset of control (using the tangent method) between joints and endpoint/hand (Fig. 2D) by performing 1-way repeated measures ANOVA (levels – shoulder, elbow, wrist, and endpoint) and found significant differences among them (*P* < 0.001). Pairwise Tukey comparisons showed that the control of joints was significantly delayed (mean ± SEM, shoulder – 161.6 ± 15.5 ms, elbow – 162.7 ± 19.2 ms, wrist – 212.9 ± 20.0 ms) as compared to the endpoint/hand (69.6 ± 3.5 ms, Adj. *P* < 0.001 for each comparison). Among the joints, the activation of the wrist was significantly delayed with respect to the shoulder (Adj. *P* = 0.043) and elbow (Adj. *P* = 0.050) joints, although there was no significant difference between the activation of shoulder and elbow joints (Adj. *P* = 0.99).

### Trajectory control of joints and endpoint during whole arm reach

We wanted to substantiate the extent of control among joints, as seen in the previous section, towards the service of an average trajectory. We performed a 2-way repeated measures ANOVA on the zero-crossings rate (*ZCR*, three factors – shoulder, elbow, and wrist; two levels – goal and fixation) and found significant differences between goal and fixation conditions (*P* < 0.001) and across the three joints (*P* = 0.034). Pairwise comparisons showed significantly higher *ZCR* during goal movements as compared to that during fixation among shoulder (mean ± SEM, goal: 0.74 ± 0.05, fixation: 0.1 ± 0.02, *P* < 0.001), elbow (goal: 0.70 ± 0.04, fixation: 0.06 ± 0.01, *P* < 0.001) and wrist (goal: 0.54 ± 0.06, fixation: 0.1 ± 0.03, *P* < 0.001) joints.

Having considered significant trajectory control among the joints, we compared the extent of control with that of the endpoint (Fig. 3A). We performed a 1-way repeated measures ANOVA on *ZCR* (levels – shoulder, elbow, wrist, and endpoint). We found a significant difference between them (*P* < 0.001). Pairwise, the Tukey test showed that control of the endpoint (mean ± SEM: 1.52 ± 0.03) was significantly greater than the shoulder, elbow, and wrist joints (Adj. *P* < 0.001 each). Also, among joints, we found significant differences in *ZCR* between shoulder and wrist (Adj. *P* = 0.003), and elbow and wrist (Adj. *P* = 0.020) joints but not between shoulder and elbow (Adj. *P* = 0.911).

To further assess the time course of control, we dissected *ZCR* into three phases (Fig. 3B, E – early, M-middle, and L – late) along the course of movement. We performed a 2-way repeated measures ANOVA on the *ZCR* (three factors – shoulder, elbow, and wrist joints; three levels – early, middle, and late phases) and found that it was significantly different across different phases of movement (*P* < 0.001) and across the joints (*P* = 0.01). For the trajectories of endpoint, when compared to the middle phase (mean ± SEM: 0.40 ± 0.01), we found significantly greater *ZCR* during the early (0.59 ± 0.01, *P* < 0.001) and late (0.53 ± 0.02, *P* < 0.001) phases of movement (26). However, unlike the endpoint, we found that the joint control towards their respective average trajectories decreased along the movement progression. The pairwise Tukey test showed that the *ZCR* during the late phase was significantly less than the early (Adj. *P* < 0.001) and the middle (Adj. *P* < 0.001) phases, but no significant differences were found between the early and the middle phases (Adj. *P* = 0.141).

### The relationship of joint control with task space and null space variability

To further understand the nature of joint control, we wanted to test how the joints explored the task space and the null space. By definition, the task space variability leads to an exploration of the endpoint, while the null space variability explores the redundancy present in the system. We plotted the task space and the null space and found that the system explored significantly more of the null space (Fig. S1, mean ± SEM: 0.030 ± 0.007) than the task space (0.010 ± 0.002, *P* = 0.004) which is a signature of good joint synergy (8, 30). We also observed that both the null space (*P* = 0.008) and the task space explorations increased significantly (*P* < 0.001) during the movement.

Given that joints show some degree of online control, we wanted to investigate how joints contribute to modulating the task space. To do this, we correlated the average decorrelation measure 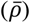 of the shoulder, elbow, and wrist joints with the task space (TS, see methods). We found that the decorrelation at the shoulder (Fig. 4A, r = 0.64, *P* = 0.010) and elbow (Fig. 4B, r = 0.72, *P* = 0.002) were significantly correlated with the task space variability. The positive correlation implied that the greater online control at joints was associated with reduced task space variability. In contrast, the wrist (Fig. 4C, r = 0.17, *P* = 0.548) was not correlated with task space variability. To understand the contribution of the wrist joint in motor control, we correlated the average decorrelation measure with the redundant null space. We found that the decorrelation at wrist (Fig. 4F, r = 0.67, *P* = 0.007) and elbow (Fig. 4E, r = 0.60, *P* = 0.018) were significantly correlated with the null space while it was weakly correlated at the shoulder (Fig. 4D, r = 0.48, *P* = 0.067).

### Leading and lagging joints during the movement

From the previous analysis, we found that the major contributors to the endpoint were the shoulder and elbow joints, and that the wrist was majorly associated with the exploration of redundancy. It was not clear whether both shoulder and elbow joints equally contributed towards the production of movement or whether they employed a leading joint strategy (13, 17). To address this question, we plotted the contribution of each joint to the endpoint velocity, in order to identify the leading and the lagging joints. Firstly, the model described in Eq. 13, was able to sufficiently predict the actual endpoint velocity (ratio of predicted velocity to actual velocity (median [Q1, Q3]): 0.89 [0.83, 0.91]). For movements directed towards 0º (Fig. 5A), we found that the shoulder joint (mean ± SEM: 97.2 ± 2.5 %, *P* <0.001) dominantly contributed towards endpoint velocity as compared to elbow (-5.1 ± 2.2 %, *P* <0.001) and wrist (6.4 ± 0.6 %, *P* <0.001). We found that the average decorrelation 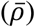 was significantly greater in shoulder (Fig. 5B, mean ± SEM: 0.81 ± 0.03) than the elbow (0.90 ± 0.02, *P* < 0.001), and the onset of online control was significantly earlier in shoulder than the elbow (mean ± SEM: -58.0 ± 27.0 ms, *P* = 0.025), and shoulder had significantly greater *ZCR* (Fig. 5C, mean ± SEM: 0.76 ± 0.06) than the elbow joint (0.46 ± 0.06, *P* < 0.001).

However, for movements directed towards 45º (Fig. 5D), the strategy switched over, and the elbow (mean ± SEM: 99.8 ± 0.03 %, *P* <0.001) dominantly contributed towards endpoint velocity than the shoulder (-4.1 ± 2.1 %, *P* <0.001) and wrist joints (4.2 ± 1.0 %, *P* <0.001). Interestingly, the average decorrelation was significantly greater in elbow (Fig. 5E, mean ± SEM: 0.84 ± 0.02) than the shoulder (0.91 ± 0.01, *P* < 0.001), and the onset of online control was much earlier in elbow than the shoulder (mean ± SEM: -86.6 ± 22.8 ms, *P* < 0.001), and elbow had significantly more *ZCR* (Fig. 5F, mean ± SEM: 0.67 ± 0.07) than the shoulder (0.37 ± 0.05, *P* < 0.001).

Taken together, we found that the shoulder was the leading joint along the directions – 0º, 135º, 180º and 315º (Fig. S2A, *P* < 0.001 each), while elbow was the leading joint along directions – 45º, 90º, 225º and 270º (*P* < 0.001 each). We averaged the velocity contribution of the leading and the lagging joints across all the directions of movement (Fig. 5G) respectively and found that the leading joint on an average contributed to 112.6 ± 2.6 %, (mean ± SEM, *P* < 0.001) of the endpoint velocity while the lagging joint impeded the movement (mean ± SEM: -18.2 ± 2.7 %, *P* <0.001). Consequently, we found that the leading joint had significantly greater decorrelation (Figs. 5H and S2B, mean ± SEM: 0.77 ± 0.02) than the lagging joint (0.82 ± 0.02, *P* < 0.001). the online control was initiated significantly earlier in the leading joint when compared to the lagging joint (Fig. 6, mean ± SEM: -60.7 ± 10.7 ms, *P* < 0.001) and *ZCR* was significantly higher in the leading joint (Figs. 5I and S2C, mean ± SEM: 0.80 ± 0.05) than the lagging joint (0.64 ± 0.04, *P* < 0.001). The wrist was never the leading joint (Figs. 5A, 5D, 5G, and S2A) and contributed to less than 6% of total endpoint velocity (5.1 ± 0.4 %, *P* = 0.02). Also, there were no systematic associations of decorrelation and *ZCR* measures across subjects and different directions of movement. However, the wrist was either lagging or assisting the leading joint but never had greater decorrelation (Figs. S2D and S2E) or greater *ZCR* (Figs. S2F and S2G) than that of the leading joint.

**Figure 6:**
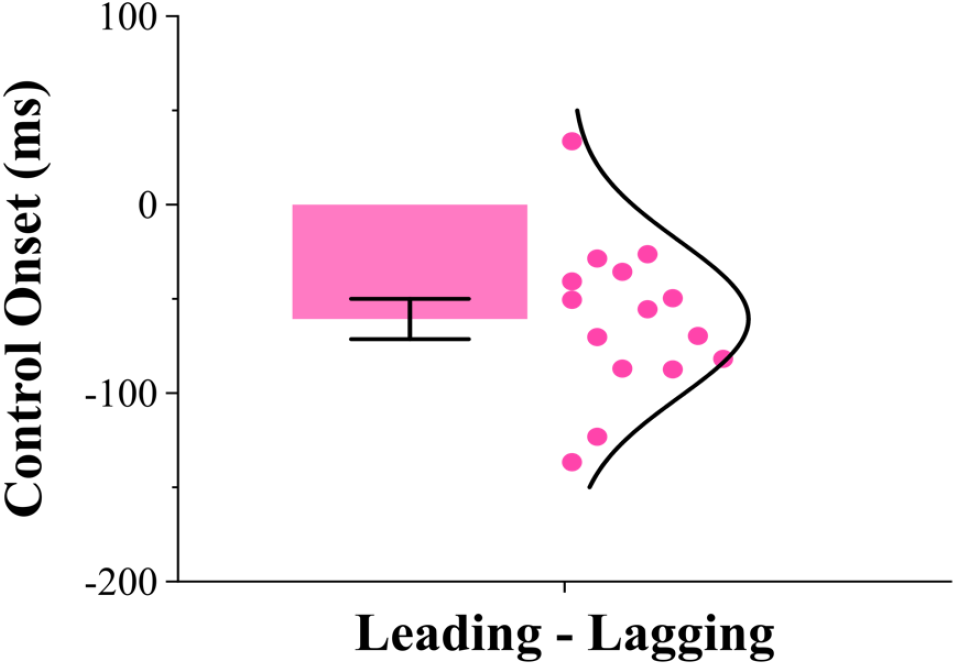
Difference between the time of onset of control of leading and lagging joints. We found that the control of leading joint preceded the lagging joint significantly by 60.7 ± 10.7 ms (mean ± SEM, P < 0.001).

## Discussion

We found differential signatures of online control among the three participating joints, potentially in the service of their respective average joint trajectories. We associated our measures of control with the pre-existing frameworks on multijointed movements such as the uncontrolled manifold hypothesis and the leading joint hypothesis and found that each of the joints adopted a unique objective in the production of reaching movements. We unravel a plausible mechanism of how the joints and the endpoint, could be planned and controlled during simple reaching movements.

### Signatures of joint control

In literature, online control or trajectory corrections have been extensively studied as the lack of predictability of future trajectory from early markers (using R^2^), in the context of endpoint during isometric force production task (18) and reaching movements (19, 21, 23), but the role of joints have largely been ignored. One such study on joints, proposed the use of canonical correlation analysis to account for multijointed movements where a high-dimensional joint posture at any time point was compared with the joint posture at the end of movement and showed differential joint control strategy between healthy and ischemic conditions (24). We performed this analysis on the three-dimensional joint trajectories (rotations at the shoulder, elbow, and wrist joints), and two-dimensional endpoint trajectories (X and Y co-ordinates of hand) with a critical difference that, given an initial joint or endpoint posture, we wanted to see how the respective trajectories evolved during the movement, and we found signatures of online control of joints perhaps to a lesser extent than the endpoint (Fig. 2A). A potential problem with this approach though is the use of linear combination of joint angles in a highly non-linear biomechanical system of the arm, and it did not provide granularity to distinguish the roles of individual joints during the movement.

To address the above problems, we performed Spearman’s correlation analysis on individual joint trajectories and on the endpoint and found qualitatively similar results between the control of the joints and the endpoint, but interestingly online control among joints were not merely a scaled down version of the control at endpoint, rather we found differential control signatures along with their respective onsets of kinematic control especially between the control of shoulder and elbow joints versus the control of the wrist (Fig. 2B and 2D). The trajectories of the shoulder and elbow experienced significantly greater and faster online control than the wrist joint by ∼50 ms. These kinematic signatures are in broad agreement with the leading joint hypothesis where it was observed that the shoulder or elbow generated movement of the limb with most of the active muscle torque as compared to the wrist joint (2, 13, 14). Additionally, the decorrelation measure varied significantly across different directions of movement (Fig. S2) indicating the necessity to control joints differentially depending on the task, similar to a mechanism described by LJH (13, 16, 17). Further, these differences in online control were not merely due to computational noise, but meaningful, as they were tightly correlated with their respective joint dispersion measures (Fig. 2B). These were also not due to noise from data acquisition as they were significantly different during the movement as compared to fixating at the center.

### Trajectory control of joints

Several studies have found that the CNS does not follow a unique joint trajectory but rather explores multiple possible joint configurations in the service of the endpoint (3, 4, 6, 10). Be that as it may, small angular deviations at more proximal joints is expected to have large effect on the location of endpoint (3), and ought to be controlled tighter than the distal joints. The zero crossings rate (*ZCR*) on the entire duration as well as its trifurcation into early, middle and late phases, was previously shown to be a reliable measure to capture corrections towards an average trajectory between goal and non-goal conditions (26). We extended this technique to analyze trajectory control of the three participating joints and found that proximal joints such as shoulder and elbow, experienced significantly greater zero crossings than the wrist joint (Fig. 3A), which is consistent with the leading joint hypothesis (2) but in the kinematic space. Interestingly, the *ZCR* at all the three joints faded out as movements approached the late phase (Fig. 3B) where it’s been noticed that the requirements of control on the endpoint becomes crucial during this phase (1, 4, 11, 37).

### Control strategy at joints vs endpoint

Despite signatures of online trajectory control at individual joint, the extent of control among joints, however, was significantly less and slower than that at the endpoint (Figs. 2 and 3). We surmise that this distinction could be due to multiple reasons. Firstly, the CNS had access to the cursor location on the screen as visual feedback of the endpoint and could integrate that with the delayed proprioceptive signals to control the endpoint while it could only tap into the latter arising from the arm to control joint movements (38). We know from several studies that visual feedback is more reliable and accurate than the proprioceptive signals (39) and that it dominates perception. But in the absence of vision, the system relies on proprioception (40, 41). Secondly, the task defined on the endpoint was implicitly shared among the participating joints (3). In a previous study, we had observed significantly greater online control and trajectory correction in the context of flexion-extension movements among fingers when the task was defined on the metacarpophalangeal joint (26, 33). But in the context of whole arm reaching, finding differential signatures of control among joints even when the task was at the endpoint (Figs. 2 and 3), hints at a dual control model where a fraction of the task could be encoded or resolved kinematically in joint co-ordinates similar to suggestions from LJH (2, 15), along with a kinematic controller implemented simultaneously at the endpoint. Interestingly, the extent of kinematic control among joints was high during the early and middle phases and decreased significantly d of movement (Fig. 3B), unlike the endpoint where greater control was observed during the early and the late phases of movement (26). While its well known that the end of movement accuracy at the endpoint is critical during tasks like these (1, 27), greater kinematic control of joints and endpoint during the early phase makes it easier for the CNS to restrict the arm from moving away from a planned trajectory.

### Is the null space an uncontrolled manifold?

The uncontrolled manifold hypothesis (UCM) states that the redundancy among joints is resolved by enforcing control over subspaces of joint coordination in the service of endpoint (5, 7, 30, 42). According to UCM, the projection of joint variability along the null space does not affect the location of endpoint, and is called the uncontrolled manifold, where it is hypothesized that CNS does not control the variability along this subspace (5). In contrast, the projection of joint variability along the task space, perpendicular to null space, is expected to produce changes in the endpoint and is hypothesized to be tightly controlled. In this regard, we observed significantly greater exploration along the null space when compared to the task space (Fig. S1), similar to previous studies (5, 30), suggesting the presence of a robust synergy among the participating joints, consistent with the UCM hypothesis.

We came up with an independent measure to quantify the online control among joints and specifically hypothesised, based on UCM, that any online control among joints, would be associated with regulating the task space and not the null space. However, we found that the online control only at shoulder and elbow joints was associated with the task space significantly (Figs. 4A – 4C), while surprisingly, online control at the wrist joint was associated with the null space (Fig. 4F). Further, the association of online control at wrist joint with the null space was robust across different directions where shoulder or elbow was the leading joint (although not shown in results). In general terms, greater online control was associated with reduced variability along both the subspaces (Fig. 4). These observations lead us to propose a modification of the critical assumption underlying UCM on null space, from an “uncontrolled manifold” to a “selectively controlled manifold”, where a few joints, mostly the distal ones, are utilized to explore or exploit redundancy. Our results are in broad agreement with a study which was able to successfully reinforce participants to explore the null space to modify the wrist joint either towards flexion or extension directions in order to maintain the required endpoint accuracy (43). They also are consistent with earlier observations that redundancy can help in motor learning (36) and could be potential used in the context of rehabilitation, where patients could be reinforced to regain movements by actively exploring the null space.

### Kinematic control of joints and the leading joint hypothesis

According to the leading joint hypothesis, the control is mediated at leading joints such as shoulder or elbow, to drive the movement of the arm by producing active muscle torque while the other lagging joints utilize interaction torques for their movement and cater to the needs of endpoint (2). The proposed control variable in LJH was in terms of the torque applied at individual joints, but it was unclear as to how reliably the system directly or indirectly tracks the torque signals at joints during the movement (44). Based on the results of differential kinematic control among joints along with its systematic variation across different movement directions (Fig. 5), which matches the predictions of LJH, suggests that the control variable could be in terms of the kinematic trajectories of the joints, and more prominently on the kinematics of the leading joint. The CNS receives proprioceptive signals about joint position and rotation (45) from various contractile sources such as muscle spindles and Golgi tendon organs, and non-contractile sources such as Type I – III receptors in the joint capsule and the cutaneous receptors from the skin (44, 46–50). The afferent signals from muscles, joints and skin integrate as they ascend upstream at cuneate nucleus (51) and thalamic cells (52) to provide information about the whole range of joint movement. Hence, it would not be surprising that the system would make use of these signals to control joint trajectories during the movement.

In the context of the LJH, the leading joint need not always be the proximal joint, rather it switches between shoulder and elbow (13, 16, 17). For a task involving drawing on a horizontal plane, it was observed that the shoulder was leading for most directions except for when the line was drawn along 45º – 225º direction with respect to the X – axis, where the elbow took over the role of the leading joint (17). Along with 45º – 225º direction (Fig. 5), we also found elbow to lead along the 90º – 270º directions. We observed a similar switch in our kinematic measures of joint control across these directions in line with the predictions from LJH. One critical difference among the two tasks was that we had the wrist free to rotate (flexion-extension) in our experiment while it was restricted using a splint in their experiment (17). We believe that adding an additional degree of freedom enabled a flexible control strategy (3). Nevertheless, irrespective of the direction of movement, the leading joint experienced a greater kinematic online control potentially towards its average trajectory as compared to the lagging joint (Fig. 5G – 5I).

Another aspect of kinematic control of joints we observed, was related to the notion of proximal to distal sequencing, which refers to the sequential recruitment of proximal to distal joints during the execution of specific kinds of movements (53, 54). Several studies have indicated such a hierarchy in recruitment using kinematic variables such as maximum linear and angular velocities, EMG onsets, and even with the recruitment of cells in the primary motor area (55–58). Our findings are in broad agreement with this recruitment pattern, particularly with respect to the onset of kinematic online control where we found online control of shoulder and elbow preceded the wrist joint on an average by ∼50 ms (Fig. 2D). But looking deeper across different directions of movement, we extend this strategy to simple reaching movements by calling it “leading to lagging sequencing” where the kinematic control of the leading joint preceded the lagging joint by ∼ 60 ms (Fig. 6).

In conclusion, along with endpoint control, we found evidence for online control of joints potentially towards their respective average trajectories. We found that while some joints such as shoulder or elbow, took over the role of leading/generating the reaching movements, the other distal joints such as the wrist was responsible for regulating the exploration of redundancy in the system. Thus, the problem of joint redundancy could potentially be resolved by allocating different objectives to different participating joints during the movement. However, further experiments are needed to extend these results to 3-D reaching movements.

## Ethical Statement

Experiments discussed in this manuscript have been ethically approved and all the participants provided their consent for the experiments. (Check methods section for further details).

## Data Availability

Original data and code supporting the findings of this study will be made available to editors and reviewers upon request during the peer review process and to readers upon publication.

## Acknowledgements

We would like to thank all the subjects who participated in this study and colleagues in the lab for their assistance during the experiments. This research was supported by the Department of Biotechnology – Indian Institute of Science Partnership Program grant (DBT– IISc, *BT/PR27952/INF/22/212/2018*), Intramural funds from IISc (Institute of Eminence grant from the Ministry of Education), and Science and Engineering Research Board grant (SERB, *CRG/2022/000553*). NC was supported by the Prime Minister’s Research Fellowship from the Ministry of Education (PMRF, *PM/MHRD-18-16074*.*03*).

## References

1. D. M. Wolpert, Z. Ghahramani, Computational principles of movement neuroscience. Nat Neurosci 3, 1212–1217 (2000).

2. N. Dounskaia, Control of human limb movements: the leading joint hypothesis and its practical applications. Exerc Sport Sci Rev 38, 201–208 (2010).

3. M. L. Latash, Synergy (Oxford University Press, 2008).

4. N. A. Bernshtein, The co-ordination and regulation of movements (Pergamon Press, 1967).

5. M. Latash, There is no motor redundancy in human movements. There is motor abundance. Motor Control 4, 259–260 (2000).

6. P. Morasso, Spatial control of arm movements. Experimental Brain Research 42 (1981).

7. I. M. Gelfand, M. L. Latash, On the problem of adequate language in motor control. Motor Control 2, 306–313 (1998).

8. M. L. Latash, Motor Synergies and the Equilibrium-Point Hypothesis. Motor Control 14, 294–322 (2010).

9. R. Bongaardt, O. G. Meijer, Bernstein’s Theory of Movement Behavior: Historical Development and Contemporary Relevance. Journal of Motor Behavior 32, 57–71 (2000).

10. M. L. Latash, J. P. Scholz, G. Schöner, Motor Control Strategies Revealed in the Structure of Motor Variability. Exercise and Sport Sciences Reviews 30, 26 (2002).

11. E. Todorov, M. I. Jordan, Optimal feedback control as a theory of motor coordination. Nature Neuroscience 5, 1226–1235 (2002).

12. Y.-K. Kim, R. N. Hinrichs, N. Dounskaia, Multicomponent Control Strategy Underlying Production of Maximal Hand Velocity During Horizontal Arm Swing. Journal of Neurophysiology 102, 2889–2899 (2009).

13. N. Dounskaia, Y. Shimansky, B. K. Ganter, M. E. Vidt, A simple joint control pattern dominates performance of unconstrained arm movements of daily living tasks. PLoS One 15, e0235813 (2020).

14. N. V. Dounskaia, S. P. Swinnen, C. B. Walter, A. J. Spaepen, S. M. P. Verschueren, Hierarchical control of different elbow-wrist coordination patterns. Exp Brain Res 121, 239–254 (1998).

15. N. Dounskaia, The internal model and the leading joint hypothesis: implications for control of multi-joint movements. Exp Brain Res 166, 1–16 (2005).

16. S. Ambike, J. P. Schmiedeler, The leading joint hypothesis for spatial reaching arm motions. Exp Brain Res 224, 591–603 (2013).

17. N. Dounskaia, C. J. Ketcham, G. E. Stelmach, Commonalities and differences in control of various drawing movements. Exp Brain Res 146, 11–25 (2002).

18. J. Gordon, C. Ghez, Trajectory control in targeted force impulses. III. Compensatory adjustments for initial errors. Exp Brain Res 67, 253–269 (1987).

19. J. Messier, J. F. Kalaska, Comparison of variability of initial kinematics and endpoints of reaching movements. Exp Brain Res 125, 139–152 (1999).

20. M. Desmurget, et al., Updating Target Location at the End of an Orienting Saccade Affects the Characteristics of Simple Point-to-Point Movements. Journal of Experimental Psychology: Human Perception and Performance 31, 1510–1536 (2005).

21. M. A. Khan, I. M. Franks, Online Versus Offline Processing of Visual Feedback in the Production of Component Submovements. Journal of Motor Behavior 35, 285–295 (2003).

22. B. A. Richardson, A. Ratneswaran, J. Lyons, R. Balasubramaniam, The time course of online trajectory corrections in memory-guided saccades. Exp Brain Res 212, 457–469 (2011).

23. S. Kuang, A. Gail, When adaptive control fails: Slow recovery of reduced rapid online control during reaching under reversed vision. Vision Research 110, 155–165 (2015).

24. M. Krüger, A. Straube, T. Eggert, The Propagation of Movement Variability in Time: A Methodological Approach for Discrete Movements with Multiple Degrees of Freedom. Frontiers in Computational Neuroscience 11 (2017).

25. R. J. Janse, et al., Conducting correlation analysis: important limitations and pitfalls. Clinical Kidney Journal 14, 2332–2337 (2021).

26. N. Chakrabhavi, V. Vasudevan, V. Skm, A. Ghosal, A. Murthy, Kinematic signatures of fast feedback trajectory control during goal-directed finger, arm, and saccadic eye movements. [Preprint] (2025). Available at: https://www.biorxiv.org/content/10.1101/2023.08.09.552575v3 [Accessed 11 June 2025].

27. D. Liu, E. Todorov, Evidence for the Flexible Sensorimotor Strategies Predicted by Optimal Feedback Control. J. Neurosci. 27, 9354–9368 (2007).

28. T. Cluff, S. H. Scott, Apparent and Actual Trajectory Control Depend on the Behavioral Context in Upper Limb Motor Tasks. J. Neurosci. 35, 12465–12476 (2015).

29. R. C. Oldfield, The assessment and analysis of handedness: The Edinburgh inventory. Neuropsychologia 9, 97–113 (1971).

30. J. P. Scholz, G. Schöner, M. L. Latash, Identifying the control structure of multijoint coordination during pistol shooting. Experimental Brain Research 135, 382–404 (2000).

31. A. P. Georgopoulos, J. F. Kalaska, R. Caminiti, J. T. Massey, On the relations between the direction of two-dimensional arm movements and cell discharge in primate motor cortex. J. Neurosci. 2, 1527–1537 (1982).

32. S. H. Scott, P. L. Gribble, K. M. Graham, D. W. Cabel, Dissociation between hand motion and population vectors from neural activity in motor cortex. Nature 413, 161–165 (2001).

33. N. Chakrabhavi, V. Skm, Wrist Posture Does Not Influence Finger Interdependence. J Appl Biomech 35, 410–417 (2019).

34. R. L. Bertrand, Lag Phase Is a Dynamic, Organized, Adaptive, and Evolvable Period That Prepares Bacteria for Cell Division. Journal of Bacteriology 201, 10.1128/jb.00697-18 (2019).

35. B. J. Smug, M. Opalek, M. Necki, D. Wloch-Salamon, Microbial lag calculator: A shiny-based application and an R package for calculating the duration of microbial lag phase. Methods in Ecology and Evolution 15, 301–307 (2024).

36. P. Singh, S. Jana, A. Ghosal, A. Murthy, Exploration of joint redundancy but not task space variability facilitates supervised motor learning. Proc Natl Acad Sci U S A 113, 14414–14419 (2016).

37. C. M. Harris, D. M. Wolpert, Signal-dependent noise determines motor planning. Nature 394, 780–784 (1998).

38. F. R. Sarlegna, R. L. Sainburg, The Roles of Vision and Proprioception in the Planning of Reaching Movements. Adv Exp Med Biol 629, 317–335 (2009).

39. K. R. Boff, L. Kaufman, J. P. Thomas, Eds., Handbook of perception and human performance (Wiley, 1986).

40. M. Mon-Williams, J. P. Wann, M. Jenkinson, K. Rushton, Synaesthesia in the normal limb. Proc Biol Sci 264, 1007–1010 (1997).

41. R. J. van Beers, D. M. Wolpert, P. Haggard, When Feeling Is More Important Than Seeing in Sensorimotor Adaptation. Current Biology 12, 834–837 (2002).

42. J. P. Scholz, G. Schöner, The uncontrolled manifold concept: identifying control variables for a functional task. Exp Brain Res 126, 289–306 (1999).

43. D. M. A. Mehler, A. Reichenbach, J. Klein, J. Diedrichsen, Minimizing endpoint variability through reinforcement learning during reaching movements involving shoulder, elbow and wrist. PLoS One 12, e0180803 (2017).

44. G. Macefield, S. C. Gandevia, D. Burke, Perceptual responses to microstimulation of single afferents innervating joints, muscles and skin of the human hand. The Journal of Physiology 429, 113–129 (1990).

45. J. R. Andrews, G. L. Harrelson, K. E. Wilk, Physical Rehabilitation of the Injured Athlete: Expert Consult - Online and Print (Elsevier Health Sciences, 2012).

46. P. Grigg, B. J. Greenspan, Response of primate joint afferent neurons to mechanical stimulation of knee joint. Journal of Neurophysiology 40, 1–8 (1977).

47. V. G. Macefield, L. Norcliffe-Kaufmann, J. Gutiérrez, F. B. Axelrod, H. Kaufmann, Can loss of muscle spindle afferents explain the ataxic gait in Riley–Day syndrome? Brain 134, 3198–3208 (2011).

48. V. G. Macefield, The roles of mechanoreceptors in muscle and skin in human proprioception. Current Opinion in Physiology 21, 48–56 (2021).

49. U. Proske, A reassessment of the role of joint receptors in human position sense. Exp Brain Res 241, 943–949 (2023).

50. U. Proske, Joint receptors play a role in position sense after all! The Journal of Physiology 602, 3609–3612 (2024).

51. J. Millar, Convergence of joint, cutaneous and muscle afferents onto cuneate neurones in the cat. Brain Research 175, 347–350 (1979).

52. V. B. Mountcastle, G. F. Poggio, G. Werner, The relation of thalamic cell response to peripheral stimuli varied over an intensive continuum. Journal of Neurophysiology 26, 807–834 (1963).

53. F. Danion, M. L. Latash, Motor Control: Theories, Experiments, and Applications (Oxford University Press, 2011).

54. B. Serrien, J.-P. Baeyens, The proximal-to-distal sequence in upper-limb motions on multiple levels and time scales. Human Movement Science 55, 156–171 (2017).

55. H. J. Jöris, A. J. van Muyen, G. J. van Ingen Schenau, H. C. Kemper, Force, velocity and energy flow during the overarm throw in female handball players. J Biomech 18, 409–414 (1985).

56. J. T. Murphy, Y. C. Wong, H. C. Kwan, Sequential activation of neurons in primate motor cortex during unrestrained forelimb movement. Journal of Neurophysiology 53, 435–445 (1985).

57. S. Furuya, H. Kinoshita, Roles of proximal-to-distal sequential organization of the upper limb segments in striking the keys by expert pianists. Neurosci Lett 421, 264– 269 (2007).

58. S. Furuya, H. Kinoshita, Expertise-dependent modulation of muscular and nonmuscular torques in multi-joint arm movements during piano keystroke. Neuroscience 156, 390–402 (2008).

