## Supporting Information (SI) for "Differential kinematic control and co-ordination among redundant joints during whole arm reaching movements"

**This PDF file includes:**

1. SI Methods
2. SI Figures – Figs. S1 and S2

### SI Methods

Expansion of the Jacobian matrix ($J$), as given in Eq. 6 of the main text –

$$J = \begin{bmatrix} \frac{\partial x}{\partial \theta_1} & \frac{\partial x}{\partial \theta_2} & \frac{\partial x}{\partial \theta_3} \\ \frac{\partial y}{\partial \theta_1} & \frac{\partial y}{\partial \theta_2} & \frac{\partial y}{\partial \theta_3} \end{bmatrix}$$

where, the partial derivatives were expressed as the following,

$$\frac{\partial x}{\partial \theta_1} = -l_1 \sin(\theta_1) - l_2 \sin(\theta_1 + \theta_2) - l_3 \sin(\theta_1 + \theta_2 + \theta_3)$$

$$\frac{\partial x}{\partial \theta_2} = -l_2 \sin(\theta_1 + \theta_2) - l_3 \sin(\theta_1 + \theta_2 + \theta_3)$$

$$\frac{\partial x}{\partial \theta_3} = -l_3 \sin(\theta_1 + \theta_2 + \theta_3)$$

$$\frac{\partial y}{\partial \theta_1} = l_1 \cos(\theta_1) + l_2 \cos(\theta_1 + \theta_2) + l_3 \cos(\theta_1 + \theta_2 + \theta_3)$$

$$\frac{\partial y}{\partial \theta_2} = l_2 \cos(\theta_1 + \theta_2) + l_3 \cos(\theta_1 + \theta_2 + \theta_3)$$

$$\frac{\partial y}{\partial \theta_3} = l_3 \cos(\theta_1 + \theta_2 + \theta_3)$$

*Note:* $l_1$, $l_2$, and $l_3$ were the link lengths of upper arm, forearm, and hand respectively. Further, $\theta_1$, $\theta_2$, and $\theta_3$ were the horizontal adduction – abduction of the shoulder, flexion – extension of the elbow, and flexion – extension of the wrist respectively.

### SI Figures

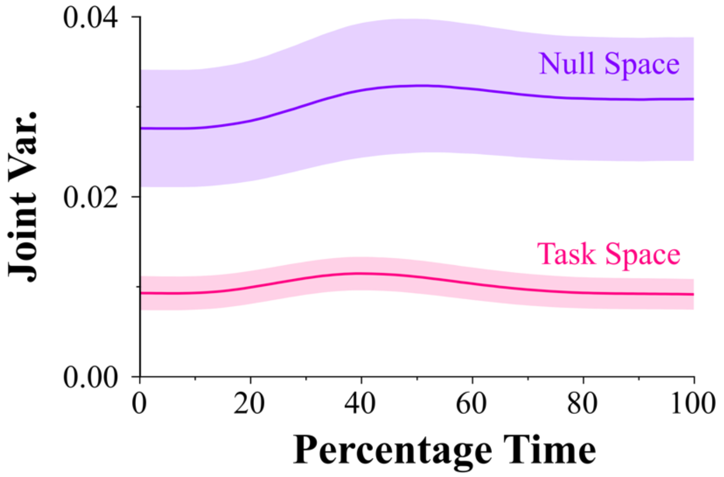


***<u>Figure S1:</u>* Plot of joint exploration along the null space (purple) and task space (pink).** *We found significantly greater joint exploration of the null space (mean ± SEM) as compared to the task space (P = 0.004). Both null space and task space exploration increased during the movement and is broadly in agreement with the previous study* (1).

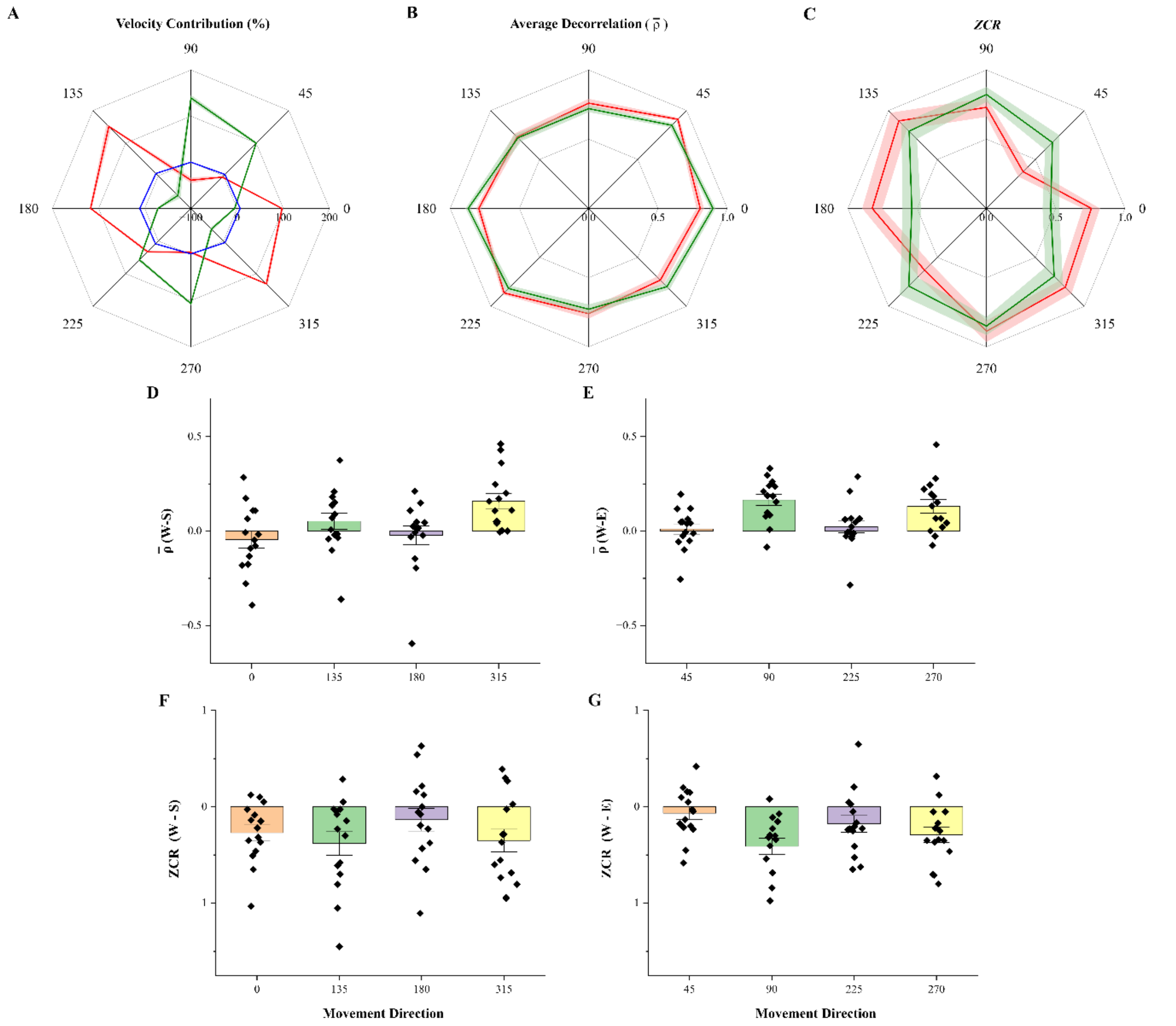


***<u>Figure S2:</u>* Plot of velocity contribution, average decorrelation ($\bar{\rho}$), and ZCR across different directions of movement (mean ± SEM).** *A) Velocity contribution of shoulder (red), elbow (green), and wrist (blue) joints towards the velocity of endpoint (hand). It could be noticed that movements were performed as per LJH* (2, 3), *where, shoulder was the leading joint for movement directions - 0º, 135º, 180º and 315º, while elbow was the leading joint for movement directions - 45º, 90º, 225º and 270º. Wrist was never the leading joint. B) Average decorrelation ($\bar{\rho}$) for shoulder (red) and elbow (green) broadly show a similar switch over like the LJH across different directions of movement. Note: Lower value indicates greater online control. C) ZCR of shoulder (red) and elbow (green) broadly show a similar switch over like the LJH. D-G) The average decorrelation and ZCR of the wrist joint (W) compared to shoulder (S) and elbow (E). When dissected into different movement directions, we did not find a systematic relationship between our kinematic measures of wrist joint with the LJH. However, we found that the average correlation co-efficient of wrist joint was never significant less (P>0.05 for all the conditions) than that of the leading joint, either when the shoulder (D) or elbow (E) was leading. But, the leading joint had significantly greater decorrelation than the wrist joint for movements along 90º (P < 0.001), 270º (P = 0.001), and 315º (P < 0.001). Similarly, the ZCR of the wrist joint was never significantly greater (P> 0.05 for all the conditions) than that of the leading joint, either when the shoulder (G) or elbow (F) was leading. On the contrary, the leading joint had significantly greater ZCR for movements along 0º (P = 0.002), 90º (P < 0.001), 135º (P = 0.004), 225º (P = 0.03), 270º (P = 0.001), and 315º (P = 0.005).*

***References –***


1. J. P. Scholz, G. Schöner, M. L. Latash, Identifying the control structure of multijoint coordination during pistol shooting. *Experimental Brain Research* **135**, 382–404 (2000).

2. N. Dounskaia, Control of human limb movements: the leading joint hypothesis and its practical applications. *Exerc Sport Sci Rev* **38**, 201–208 (2010).

3. N. Dounskaia, Y. Shimansky, B. K. Ganter, M. E. Vidt, A simple joint control pattern dominates performance of unconstrained arm movements of daily living tasks. *PLoS One* **15**, e0235813 (2020).
